# *E. coli* division machinery drives cocci development inside host cells

**DOI:** 10.1101/2024.04.08.588611

**Authors:** Alaska Pokhrel, Ariana Costas, Matthew Pittorino, Iain Duggin, Bill Söderström

## Abstract

*Escherichia coli* is arguably one of the most studied bacterial model systems in modern biology. Under normal laboratory conditions *E. coli* adopts its characteristic rod-shape. However, during stress conditions *E. coli* has been shown to undergo conditional morphology changes to inhibit division and grow into highly elongated forms. Here, on the other end of the morphology spectra, using an *in-vitro* infection model system combined with advanced imaging we show uropathogenic *E. coli* rods dividing to form and proliferate as cocci inside human bladder epithelial cells. In these intracellular bacterial communities, the frequency of cell division outpaced the rate of cell growth, resulting in smaller cocci cells. This mechanism was guided by an active FtsZ-governed division machinery, directed to midcell by division-site placement systems. These results show how a previously uncharacterised level of morphological plasticity occurs in bacteria with traditionally well-defined rod shape.

## Introduction

Intracellular bacterial division and proliferation is essential for pathogenic bacteria to extend and maximize intracellular colonization during infection in disease settings[1]. Division in the rod-shaped model bacterium *Escherichia coli* has been well mapped in laboratory settings, and is known to be carried out by a dynamic macromolecular complex led by the highly conserved FtsZ protein [2]. FtsZ organises an assembly of about 30 different proteins, which together coordinate the remodelling of the septum leading up to septation [3]. Guided by spatial regulatory systems (*e.g.*, the Min system [4] and Nucleoid occlusion [5]), FtsZ is the first essential protein that localises to midcell where it polymerizes into filaments forming an intermediate structure that resembles a ring, commonly referred to as the “Z-ring” [6]. Accumulation of FtsZ at the midcell also initiates the recruitment of other essential division proteins, of which FtsN is the last one to be recruited to the division site [7]. The arrival of FtsN to the thus far preassembled divisome complex triggers the initiation of septal cell wall synthesis resulting in inward membrane invagination, septum formation and eventually cell separation[8].

Intracellular proliferation of *E. coli* within a pathological setting is less well understood but what is clear is that during unfavourable environmental conditions, these bacteria are readily equipped to modulate their growth and division as a survival strategy [9]. One striking example of this is the ability of rod-shaped bacteria to adopt both cocci and filamentous morphologies during urinary tract infections (UTI) [10]. UTI are one of the strongest drivers of antimicrobial resistance globally [11], where Uropathogenic *E. coli* (UPEC) accounts for approximately 75-80% of all reported cases [12].

The UPEC infection cycle is largely understood in broad strokes [13], but surprisingly few molecular details of the intracellular lifecycle of UPEC during UTI are currently known. UTI is generally initiated with bacteria being internalized by superficial epithelial cells in the host bladder, where UPEC subsequently multiplies within the cytoplasm to form densely packed colonies known as intracellular bacterial communities (IBC) [14]. It has previously been observed that UPEC cells within IBCs in UTI murine models appear as cocci with cell lengths measuring around 1μm or less, and it was hypothesised that these cells simply slowly “shrank” or “transitioned” into cocci [15]. Nutrient depletion was not considered a growth limiting factor as the bacterial biomass increased over time, with generation times exceeding 60 min inside the bladder cells [16]. In a remarkable turn of morphology regulation, during the final step of the infection cycle, UPEC forms filaments measuring hundreds of micrometres in length that flux out of the host cells into the bladder lumen [14a]. These filaments would subsequently revert back to rods prior to invading surrounding cells and initiating a new round of infection [15a].

While UPEC pathogenesis during UTI is quite well understood from a clinical perspective[17], single cell division dynamics of UPEC inside the host cells has remained largely unexplored. Here, we followed the dynamics of bacterial proliferation and visualised subcellular protein localisation inside cultured human bladder cells. This study shows that the bacterial cell division machinery, and its regulatory systems are key players driving cocci development and proliferation inside host cells.

## Results

### Rods to cocci transitions during infection

To follow intracellular UPEC proliferation over time, we used an UTI *in-vitro* infection model where a layer of immortalized human epithelial PD07i bladder cells is challenged with the pathogenic UPEC strain UTI89 (Figure S1) [18]. In order to visualise UTI89 proliferation inside PD07i cells, they were transformed to express a cytoplasmic marker (*i.e.*, mCherry)[19]. The bacterial division dynamics were subsequently followed inside the bladder cells using live cell fluorescence microscopy (Figure 1a). Interestingly, around 6h post infection [15a], we observed UTI89 bacilli transition into a cocci morphology with only one or a few cells present in the bladder cell cytoplasm (Figure 1b). This observation partly contrasts previous studies stating that cocci formation is only initiated in dense IBCs, albeit in a different model system at lower time resolution [15a].

**Figure 1.**
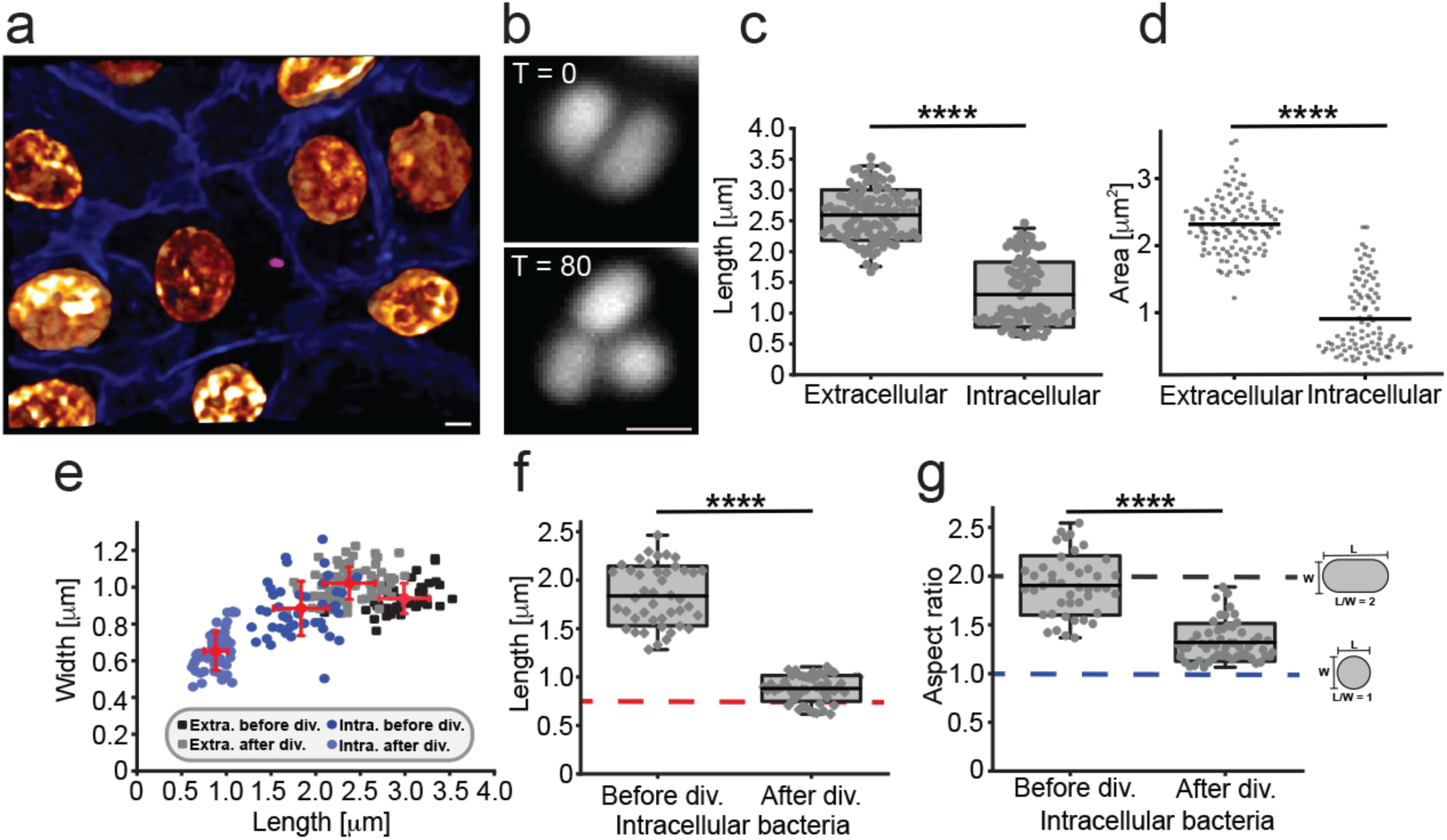
Intracellular UPEC actively divides into cocci during UTI model infection. **a**, PD07i human epithelial bladder cells (membranes blue, nuclei gold) were challenged with fluorescently labelled UTI89 (magenta). **b**, Representative images of cells before (T = 0) and after division (T = 80), note that one daughter cell has left the field of view. Times (T) are shown in minutes. **c - d**, Average cell length (**c**) and area (**d**) of extracellular and intracellular bacteria. **e**, Length vs width of extracellular (‘Extra’) and intracellular (‘Intra’) bacteria. Error bars indicate S.D. **f - g**, Lengths (**f**) and aspect ratio (**g**) of intracellular bacteria before and after division. Red dotted line in **f** represents average cocci width after division. Blue dotted line in **g** represents an aspect ratio of 1, i.e., a perfect circle. Box plots: box edge equals S.D., midline represents average, and whiskers indicate 1-99% interval. Statistical significance was determined by student’s T-test, where **** stars indicate p values less than 0.0001. Scale bars **a** = 4 μm; **b** = 1 μm.

In agreement with the previous observations, we noticed that intracellular bacterial cells were overall significantly shorter than extracellular bacteria, measuring 1.3 ± 0.53 μm (n_Intra_ = 100) and 2.59 ± 0.41 μm (n_Extra_ = 117) respectively (Figure 1c), with an overall smaller area: Area_Intra_ 0.91 ± 0.54 μm^2^ (n_Intra_ = 100) vs Area_Extra_ 2.32 ± 0.43 μm^2^ (n_Extra_ = 117) (Figure 1d). Following the division dynamics of intracellular bacteria over time (average length before division 1.84 ± 0.31μm (n = 44)), we observed UTI89 rods actively dividing into coccoid shape with average lengths of 0.88 ± 0.13 μm (n = 56) just after division (Figure 1e - f), often with an aspect ratio close to 1 (average 1.3 ± 0.19, Figure 1g). This was significantly different from the aspect ratio of extracellular cells after division 2.13 ± 0.13 (n = 76) (Figure S2). When the division dynamics was followed over multiple generations, we observed that these cocci divide into a new set of cocci, suggesting an altogether different growth strategy compared to that in planktonic cultures, where elongation outpaces division to maintain rod shape (Figure S3, Supplementary movie SM2). To determine whether this division behaviour was more broadly adapted by UPEC, we challenged bladder epithelial cells with another clinical urinary tract bacterial isolate, MS2027 [20]. We observed that both division patterns and generation times of intracellular MS2027 were indistinguishable from those of UTI89 (Figure S4). Taken together, this data clearly shows that intracellular UPEC actively divide into cocci (rather than just shrink due to the external environment) and maintain the cocci morphology through multiple generations.

### Divisome proteins drive cocci divisions in host cells

To substantiate our observation that cocci formation is driven by cell division, we followed divisome dynamics in UTI89 expressing fluorescent division markers (FtsZ-mCitrine and/or mCitrine-FtsN) during intracellular divisions (Figure 2a - b and 2d). Using FtsZ accumulation at midcell as a marker for on-going division, cocci division times ranged from roughly 50 minutes up to more than two hours (Mean: 127 ± 75 min, n = 49), and beyond 6 hours in the extreme case (Figure 2a-c). Overall, the observed apparent division times were far greater than the ∼ 20 min generation times for planktonic cultures in rich media (Figure S5). When following intracellular cocci proliferation over multiple generations, we observed that the average division time since previous division increased: 2^nd^ division average time 195 ± 81 min (n = 13) and 3^rd^ division average time 283 ± 15 min (n = 3, note only few cells could be followed over multiple generations due to the challenging 3D imaging environment) (Figure 2c). This was further confirmed by following intracellular mCitrine-FtsN dynamics in cocci cells, which indicated similar division times (Figure 2d). Similarly, FtsZ-mCitrine localization patterns and division dynamics in the MS2027 strain mirrored those in UTI89 (Figure S4).

**Figure 2.**
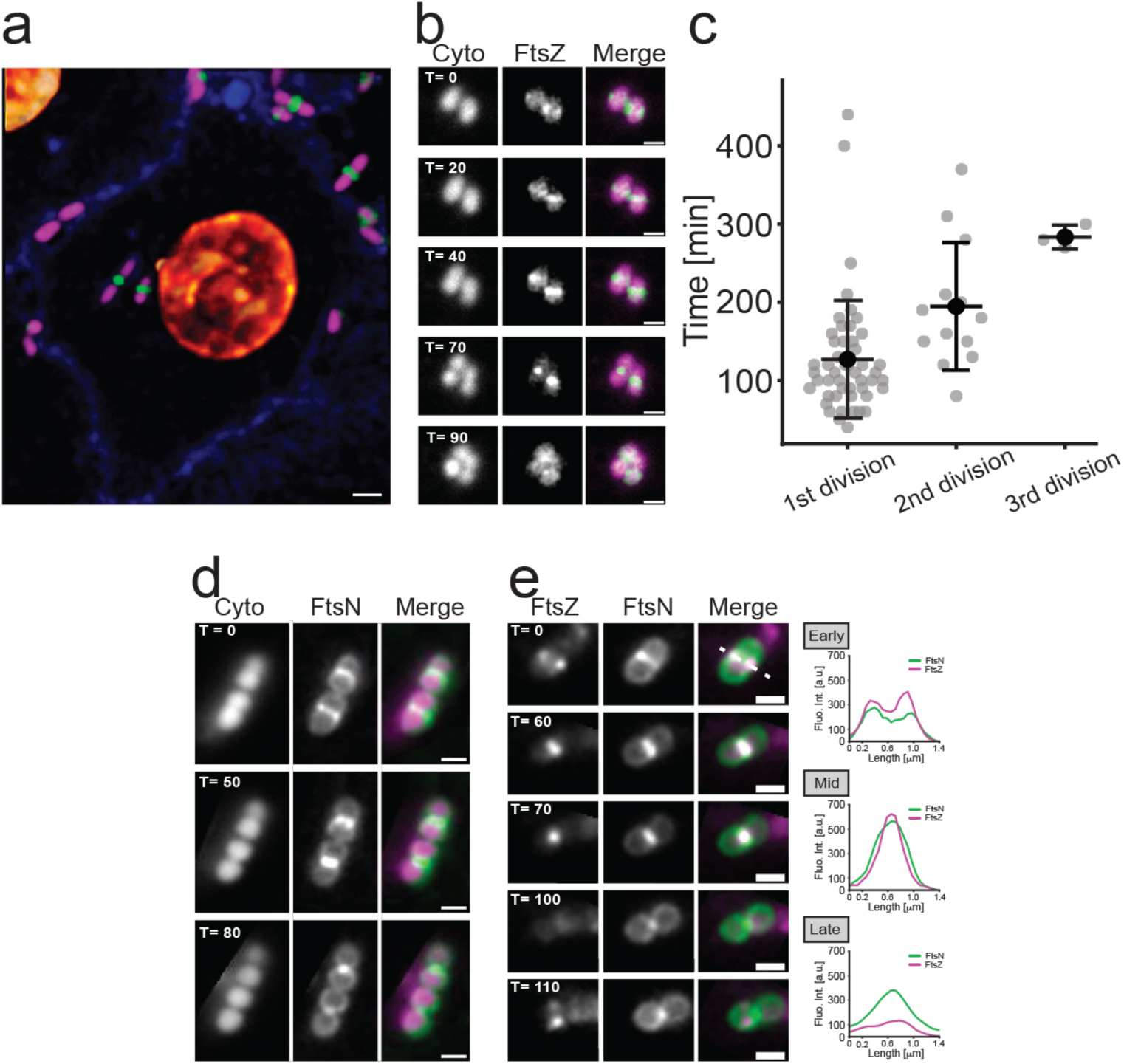
Intracellular UPEC rod-to-cocci transitions are governed by the divisome. **a**, Representative image of intracellular UTI89 expressing a division marker (FtsZ-mCitrine) with mCherry as cytoplasmic marker (bladder cell membranes in blue and nuclei in gold). **b**, Still images of a time-lapse sequence showing FtsZ-mCitrine driven cocci divisions (Supplementary movie SM3). **c**, Average division times of intracellular bacteria over multiple generations. Midline represents average, and whiskers indicate S.D. **d**, mCitrine-FtsN midcell constriction during cocci divisions (Supplementary movie SM4). **e**, FtsZ-mCherry and mCitrine-FtsN dynamics during cocci divisions. Fluorescence intensity plots highlight a larger apparent radius of FtsN (Mid, T = 70) and that it remains longer at septum than FtsZ (Late, T = 100). Times (T) are shown in minutes. Scale bars **a** = 2 μm; **b**, **d** & **e** = 1 μm

FtsZ and FtsN (the first and the last essential division protein to assemble at division sites, respectively) localization dynamics followed the same constriction patterns in cocci cells as in rods, where FtsZ-mCherry had apparent smaller radii compared to mCitrine-FtsN during constriction (Figure 2e, “mid”) [21], with FtsZ-mCherry disassembling from the division site prior to mCitrine-FtsN towards the end of division (Figure 2e, Supplementary movie SM5). Based on these data, we conclude that the divisome is actively involved in intracellular rods to cocci transition during infection, with key components of the divisome following similar localization dynamics as in vegetative growth [22]. Further, to investigate the correlation of genes previously implicated in regulating division and morphology with cocci transitions, we tested UTI89 strains where the genes encoding for SulA[23], YmfM[24] *and* DamX[25] were deleted. None of these deletion strains showed any observable impaired abilities to undergo morphology transitions from rod-to-cocci (Figure S6).

### Min oscillations in intracellular cocci

The above-mentioned findings presented an interesting conundrum, how does a normally rod-shaped cell know where to place the cell division machinery during cocci transition? In rod-shaped *E. coli* cells, the division machinery is guided to midcell by the FtsZ antagonistic effects of MinC [26]. A time-averaged concentration minima at midcell is generated by pole-to-pole oscillations of Min system led by the MinD protein[27]. However, simulations and experiments have suggested that short rods below a certain length (∼ 2.5 μm or less) do no longer undergo smooth MinD oscillations, but instead show stochastic pole-to-pole switching [28]. Here, we examined EYFP-MinD dynamics in intracellular cocci (Figure 3a), which revealed a concentration minimum at midcell both from pole-to-pole oscillations and from apparent circumferential movements along the cell perimeter indicating that the actual protein dynamics play only a minor role for the time-averaged subcellular localization patterns (and as a consequence for the time-averaged concentration profile) (Figure 3b–c, Supplementary movie SM6). Like the localization patterns in rod shaped cells, FtsZ-rings were observed at the EYFP-MinD concentration minima, and always in the same plane Z-rings from previous divisions, suggesting a maintained polarity also in cocci cells (Figure 3d). However, the average EYFP-MinD oscillation speed normalized to cell length was significantly lower in intracellular bacteria: 0.055 μm sec^-1^, compared to 0.094 μm sec^-1^ for extracellular bacteria (Figure 3e - f). Furthermore, in shorter cells, roughly shorter than 1.3 μm (much less than the 2.5 μm previously predicted), we didn’t observe the typical pole-to-pole oscillations but a much more complex EYFP-MinD dynamics, such as circular movements along the circumference and stochastic transitions from one pole to the other (Figure 3g).

**Figure 3.**
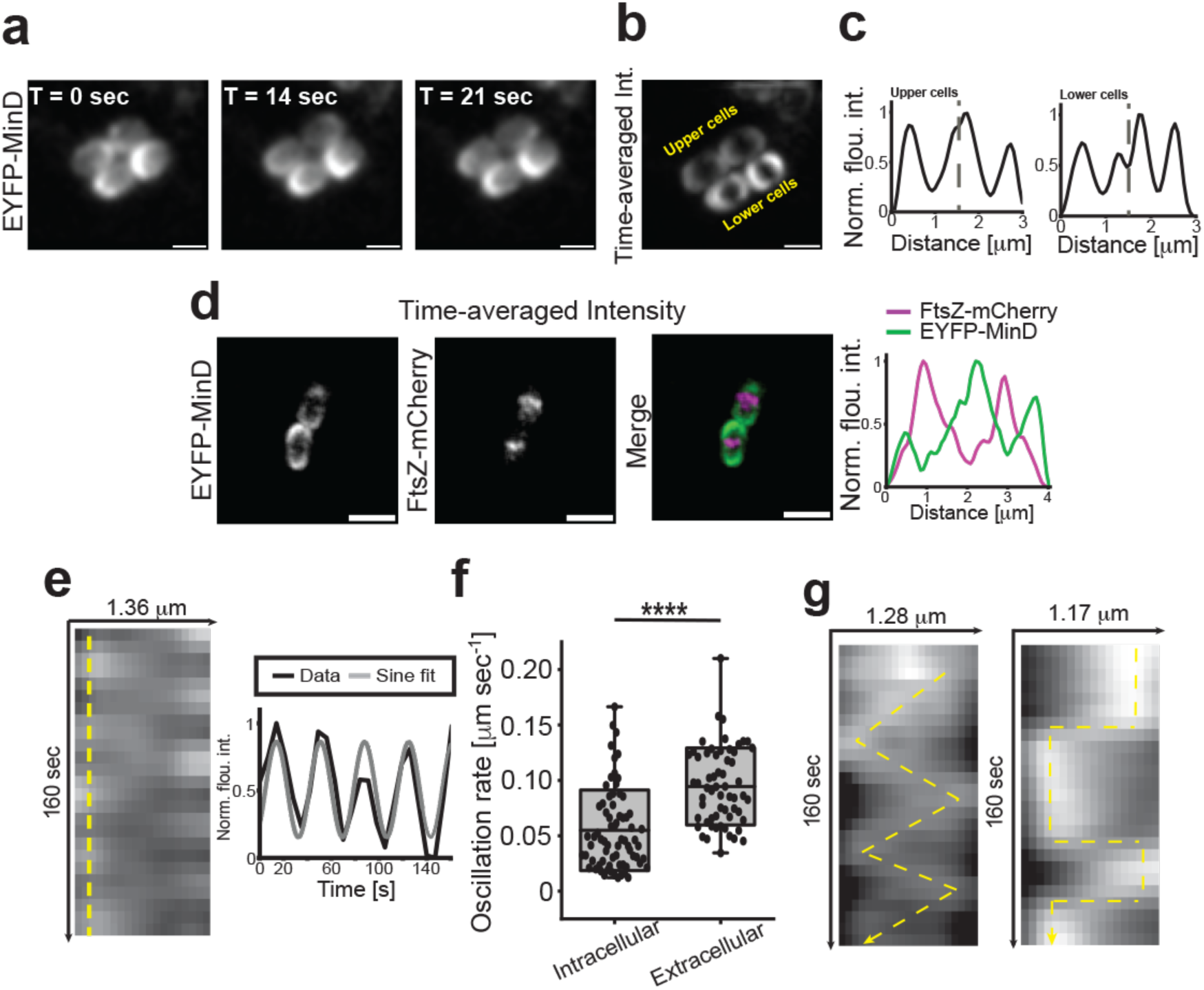
EYFP-MinD dynamics in UPEC cocci during infection. **a**, Still images from an EYFP-MinD time-lapse series showing dynamics in intracellular UTI89 cocci (Supplementary movie SM6). **b - c**, Time-averaged fluorescence intensity of EYFP-MinD. Dotted line in plots **(c)** represent cell boundaries. **d**, Time-averaged fluorescence intensity of EYFP-MinD and FtsZ-mCherry, indicating that FtsZ localize at MinD minima also in cocci. **e**, Representative kymograph of typical EYFP-MinD oscillations of a cell longer than 1.3 μm. Yellow dotted line represents where intensity plot was generated. **f**, Average EYFP-MinD oscillation rate in intracellular bacteria was 0.055 ± 0.036 μm sec^-1^ (n = 71) and in extracellular 0.095 ± 0.035 μm sec^-1^ (n = 60) cells. **g**, Typical kymographs of EYFP-MinD oscillations of cells shorter than 1.3 μm show complex non-standard pole-to-pole oscillation dynamics. Kymographs in **g** were generated from the two lower cells in Supplementary movie SM6. Scale bars **a** and **b** = 1 μm; **d** = 2 μm.

### Chromosome partitioning in intracellular cocci

Deletion of the Min system in *E. coli* is known to generate ‘mini cells’ often devoided of genetic material [29]. To investigate whether UPEC cocci had intact genetic material, and whether the nucleoid occlusion mechanism that prevents division over unsegregated chromosomes [30] was preserved during intracellular rod-to-cocci divisions, we looked at chromosome segregation inside host cells over time. When imaging intracellular UTI89 expressing a DNA marker (HupA-RFP), we never observed a cell division event where the DNA did not partition into both daughter cocci (Figure 4a). Overall, DNA partitioning in cocci was highly precise (Figure 4b), indistinguishable to what has been found in rods previously [31]. The average nucleoid length in intracellular cocci measured 0.79 ± 0.12 μm (n = 100), while extracellular cells measured 1.45 ± 0.2 μm (n = 216) (Figure 4c-d), suggesting a greater compaction of genetic material in cocci cells.

**Figure 4.**
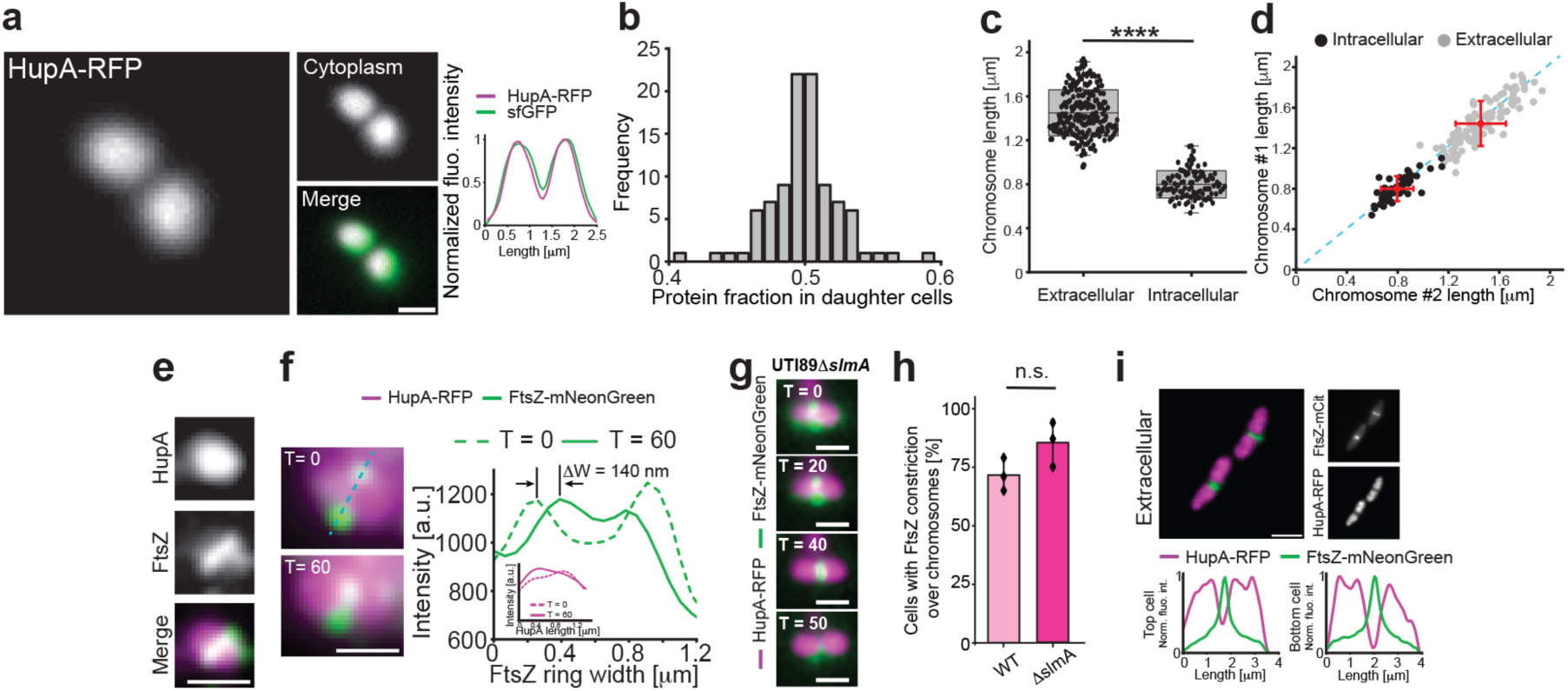
Chromosomal partitioning is maintained during intracellular cocci division. **a-b**, DNA (HupA-RFP) partitioning in cocci (msfGFP as cytoplasmic marker). DNA segregates equally into daughter cocci (n = 98). **c**, Average chromosome lengths in extracellular (n = 216) and intracellular bacteria (n = 98). **d**, Chromosomes partitions into equal size after division in both extracellular (grey) and intracellular (black) bacteria. Blue dashed line indicates linear dependence. Red dots indicate average, with x-y bars representing S.D. **e-f**, FtsZ-mNeonGreen rings assemble and, partly, constrict over undivided chromosomes (HupA-RFP) in both WT and ΔslmA cocci. Intensity plot in **f**, shows FtsZ constriction over 60 min (inset show HupA-RFP fluorescence profiles taken perpendicular to the Z-ring, with no intensity decrease at midcell over the same time interval). Cerulean dotted line indicate from where the FtsZ-mCitrine intensity plot is generated. **g**, Representative image sequence of UTI89ΔslmA expressing FtsZ-mNeonGreen and HupA-RFP showing FtsZ-ring constriction over unsegregated chromosomes. **h,** Average levels of FtsZ-mNeonGreen constriction over unsegregated chromosomes from 3 independent infections (n_cell_ = 30). **i**, FtsZ-mNeonGreen rings do not constrict over unsegregated chromosomes in extracellular rods. Statistical significance was determined by student’s T-test, where **** stars indicate p values less than 0.0001., n.s. = not significant. Box plots: box edge equals S.D., midline represents average, and whiskers indicate 1-99% interval. Times (T) are shown in minutes. Scale bars **a**, **e**, **f** and **g** = 1 μm; **i** = 2 μm.

DNA segregation in rod-shaped cells is tightly regulated by the nucleoid occlusion system [5]. In *E. coli* it is the responsibility of the SlmA protein to prevent FtsZ-ring constriction over unsegregated chromosomes, and this is achieved by complex interactions between DNA, SlmA, and the C-terminal binding domain of FtsZ [30, 32]. Given that the intracellular cocci measured only ∼ 1/3 of the length of predivisional rod-shaped cells, we wondered if full chromosomal segregation occurs prior to initiation of Z-ring constriction, or if Z-rings would partially initiate constriction prior to visible chromosome segregation. To confirm this, we looked at DNA segregation (HupA-RFP) in relation to septal constriction (FtsZ-mNeonGreen) (Figure S7). We observed that FtsZ-mNeonGreen readily formed bands over unsegregated nucleoids and appeared to initiate constriction prior to visible chromosome segregation in cocci (Figure 4e-f). The FtsZ ring radius decreased up to ∼ 140 nm (or roughly 1/3 of the diameter) prior to visible HupA-RFP intensity decrease at midcell (Figure 4f). Since Z-ring constriction was initiated over unsegregated chromosomes in WT UTI89 cocci, we also looked at Z-ring constriction in relation to DNA segregation in UTI89ι1*slmA* cocci during intracellular divisions (Figure 4g). In these cells, Z-rings initiated constriction over non-segregated nucleoids at a slightly larger degree than in WT, 85.3 (± 9.6) vs 72 (± 6.55) % of cells respectively, but with a non-statistically significant difference (p = 0.118) (Figure 4h). Overall, this behaviour was different to what was observed in extracellular cells during infection, where FtsZ-mNeonGreen constricted only when chromosomes were visibly separated (Figure 4i), even in short cells ∼ 2 - 2.5 μm, just after previous divisions (Supplementary movie SM7).

## Discussion

Here, we show that rod-shaped UPEC cells change their growth behaviour to transition into cocci in a one-step division process during intracellular proliferation in a model infection system. These morphological changes were initiated early with only one or a few UPEC cells present within the bladder cells, and the UPEC cells were seen to maintain such morphology over several generations while in an intracellular environment. While we cannot rule out a contribution of the cell elongation machinery (*e.g.*, MreB and RodZ play roles in regulating chromosome segregation and cell shape in rods [33]), intracellular rod-to-cocci divisions appears to be largely driven by an active divisome, with protein dynamics similar to those in binary fission of rod-shaped cells during vegetative growth. The cell division machinery and Min oscillations in spherically shaped *E. coli* cells have previously been investigated in mutant strains under test tube conditions [34]. However, to our knowledge, this is the first time these mechanisms have been experimentally investigated in a wild-type mode of UPEC proliferation inside host cells. Our data show FtsZ and FtsN localization patterns in intracellular cocci during constriction were indistinguishable to those in rods, but interdivision times were on average six times longer. In line with this, MinD dynamics were two times slower in cocci with pole-to-pole transitions switching from continuous oscillations to stochastic with decreasing cell size. The switching between continuous and stochastic oscillation behaviour of MinD has previously been modelled to take place in cells about 2 μm or less in length, but here we show that this phenomenon takes place in cells considerably shorter than that, roughly ∼ 1.3 μm in length. With the average MinD concentrations maintained at a minimum at midcell in cocci, the chromosome partitioning was also precisely maintained during cocci division as previously shown in rod-shaped non-pathogenic *E. coli*. The slower MinD pole-to-pole oscillations in cocci could reflect the fact that the same amount of DNA (as in rods) has to be packed into a smaller cell volume without introducing errors, which could partly explain the overall prolonged cocci division times. A somewhat surprising observation in our experiments, and in stark contrast to what has been shown in rods, was that FtsZ constriction appeared to be initiated prior to visual nucleoid segregation. While we currently don’t fully understand the reasons behind this, we speculate that SlmA has a less pronounced role during intracellular cocci divisions since efficient chromosome partition was preserved in cells lacking *slmA* altogether.

On an cellular infection level, one plausible explanation for intracellular pathogens to divide into cocci shaped cells could be to allow for a higher number of bacteria per infected bladder cell, up to two orders of magnitude more for unorganised bacilli [35], or for a greater compaction of bacteria in IBCs, where UPEC cocci have shown a survival advantage over rods within intestine in murine models[15b]. Thus, it is possible that division into cocci with longer interdivision times during infection provides intracellular survival and persistence advantages [36].

The ability to inhibit division to regulate its morphology and elongate into several hundred of micrometres in length in response to stress is another known form of UPEC plasticity observed during dispersal stages of infection in clinical settings [14a]. This switch in morphology regulation was initially attributed to the SOS system[23], although more recent studies have demonstrated filamentation in UPEC strains where the SOS regulators were absent (UTI891′*sulA*1′*ymfM*) in both murine and *in-vitro* flow models [24], suggesting a deeper level of regulation currently not well characterized. Here, we investigated the effect of deletions of the SOS-system associated genes and other components associated with infection-related filament regulation on intracellular cocci development and division dynamics. However, we did not detect apparent changes in cocci transition in any of the tested deletion strains, pointing to a different mode of cell shape regulation altogether. As cocci formation has previously been linked to increased survival during intracellular growth in mice[15b], we speculate that UPEC rod-to-cocci morphological transition are largely a result of the low nutrient availability in the hostile environment presented during intracellular life in human host cells.

## Methods

### Bacteria growth and protein production

Bacterial strains used throughout the study were routinely plated on LB agar (Difco) plates supplemented with appropriate antibiotics (Ampicillin: 100 μg ml^-1^ (Sigma), Spectinomycin 100 μg ml^-1^ (Sigma), Kanamycin 50 μg ml^-1^ (Sigma)). A single colony of respective *E. coli* strain was grown in LB media (Difco) supplemented with appropriate antibiotics at 37°C overnight without shaking. Following overnight growth, cultures were were pelleted and resuspended in 1x PBS (Bio-Rad) to an optical density (OD_600_) of 0.1. Fluorescent protein fusion production was initiated by addition of 1μM IPTG for FtsZ-mCitrine (pHC054) in UTI89 (2μM IPTG in MS2027), 5μM IPTG for EYFP-MinD (pSR-4) and HupA-RFP (pSTC011), and 15μM Rhamnose for mCitrine-FtsN (pHC004). Note that FtsZ-mNeonGreen (pMP7), FtsZ-mCherry (pMP11), msfGFP (pGI5) and mCherry (pGI6) did not require induction as they are produced under constitutive promoters. Cell viability, generation times, and cell dimensions were comparable to those of a WT strain grown under the same conditions (Figure S5). Bacterial strains and plasmids used in this study are listed in Supplementary table 1.

### Cell culture

Immortalized human epithelial bladder cells PD07i[37] were maintained in antibiotic free growth EpiLife Medium (Gibco) supplemented with growth supplement (HKGS) at 37°C in the presence of 5% CO_2_. Cells were passaged using standard trypsinization protocols upon reaching ∼ 75 - 90% confluency. To prepare for infection, cells were seeded in 35 mm glass bottom petri-dishes (IBIDI, #1.5H glass, Cat.No:81158) and grown to confluency.

### Infection model

Seeded petri-dishes were challenged with appropriate bacterial strain at an MOI of ∼ 100 (roughly corresponding to 10^7^ cells as determined by CFU counts). To facilitate increased bacterial adherence to the bladder cells, dishes were spun at 500 rpm for 3 minutes, before incubation at 37°C (5% CO_2_) with orbital agitation (50 rpm) for 15 minutes. Media containing bacteria was removed and the petri-dishes were replenished with fresh pre-warmed EpiLife media supplemented with inducer as appropriate. Dishes were incubated for at least 6 hours to let bacteria invade the bladder cells and initiate intracellular divisions (which coincides with “middle IBC” maturation[15a]). Following incubation, media was removed from petri-dish and cells were washed once with 1 x PBS. The bladder cells were stained for membrane (CellBrite Steady Membrane 405) and nucleus (NucSpot Live Cell 650) for 30 minutes in pre-warmed EpiLife as per manufacturer’s instructions (Biotium, San Francisco, USA). After the staining step and two washes with 1 x PBS, cells were covered in fresh pre-warmed EpiLife (supplemented with inducer as required) and Gentamicin f.c. 50 μg ml^-1^ (Sigma) and taken for imaging. Only intracellular bacteria survive this standard Gentamicin protection assay treatment[38], and thus dividing bacteria are inside bladder cells (Supplementary movie SM1) while non-dividing bacteria are extracellular (Supplementary movie SM8). Generally, all infection experiments were run in at least three independent replicates, unless otherwise indicated.

### Imaging

For time-lapse imaging, live cells were imaged in the glass bottom petri dishes on a Nikon N-STORM v5 TiE2 (with NIS v.5.30) microscope in TIRF mode using HILO illumination, with a 100 x 1.49 NA oil objective, to increase the signal-to-noise ratio. To support live cell imaging, the microscope was equipped with a stage top environmental chamber regulating temperature (37°C) and CO_2_ levels (5% CO_2_). 405, 488, 561 and 647 nm laser lines were used to excite fluorescent probes (405: membrane dye. 488: mCitrine, mNeonGreen, EYFP. 561: mCherry. 647: nuclear dye) as appropriate. Acquisition times were 70-400 ms and image size 2048 x 2048 (pixel size 65 nm). Fluorescence emission was collected via emission filters sets for DAPI, FITC, TxRed and ‘Normal STORM(647)’. Images were captured using a sCMOS Flash 4.0 v3 (Hamamatsu) camera. To follow the bacterial cell division within live bladder cells, images were acquired every 7 - 10 minutes for at least 6 hours (Note: it was generally not possible to accurately track more than 1 division event due to the intracellular movements with bacteria transitioning in and out of focus). To follow short term EYFP-MinD dynamics, images were acquired every 5 - 7 seconds for at least 2 minutes.

For confocal live cell imaging, images were acquired on a Leica Stellaris 8 confocal microscope equipped with a 63x NA1.49 oil objective enclosed in an environmental chamber at 37°C in the presence of 5% CO_2_. Fluorophores were excited using a white laser, and emission was collected using software optimized settings in the LAS software. Image size was chosen to either 2048 x 2048 or 4096 x 4096 pixels, with individual pixel size 90 and 45 nm, respectively. The pinhole was set to 1 AU and Z-stacks were collected with step length of either 125 or 250 nm (15 - 99 images per stack, depending on step length and thickness of the bladder cell in question).

### Image postprocessing

Raw microscopy images were transferred to Fiji for processing and analysis. Time-lapse image sequences were drift-corrected using StackReg or HyperStackreg where appropriate, and background was subtracted using a rolling ball radius of 50. Images were denoised using the PureDenoise plug-in. To follow division of intracellular bacteria/cocci (Figure 1 b-g), time-lapse sequences were segmented based on cytoplasmic fluorescent protein intensity using the tracking workflow in the ilastik Fiji plug-in (version 1.4.0). Divisions were determined being over at the time point of the first image of visually separated cells. EYFP-MinD based kymographs were generated using KymoResliceWide. Time-averaged fluorescence intensity images of EYFP-MinD were generated using the standard deviation option in the Z-projection tab. Deconvolution and reconstruction of confocal Z-stacks were performed using Leica LAS.

### Data analysis and statistics

Cell length, area, aspect ratio, and chromosome length of intra- and extra-cellular cells (rods and cocci) were extracted by using the MicrobeJ [39] plug-in or from manually traced dimensions in Fiji. Intracellular UPEC expressing fluorescent protein fusions to a division protein were followed for 1 - 3 divisions in time-lapse movies to manually extract intracellular division times. Visible FtsZ-FP accumulation at midcell was considered as initiation of division, and its disassembly from midcell was considered as completed division. Generally, fluorescence intensity profiles were generated in Fiji and analysed in Origin Pro 2021 Academic (V9.8.0.200). EYFP-MinD oscillation rates per length in intra- and extra-cellular cells were determined by measuring cell lengths and extracting fluorescence profiles over time generating corresponding kymographs. Statistical analyses were performed in either Origin Pro 2021 Academic or GraphPad Prism 10 (V10.0.3). Statistical significance was datamined by student’s T-test, where **** stars indicate p values less than 0.0001, unless defined otherwise. Figures were prepared using Adobe Illustrator.

### Western blotting

A culture volume corresponding to 0.1 OD units was collected from each UPEC strain expressing the protein of interest after at least 6 hours of growth in PD07i seeded petri-dishes (Figure S5). The samples were pelleted and re-suspended in SDS-loading buffer and resolved by SDS-PAGE gel electrophoresis. Proteins of interest (FtsZ, FtsN and MinD) were transferred to nitrocellulose membranes using a semi-dry Transfer-Blot apparatus (Bio-Rad). The Nitrocellulose membranes were then blocked in 5% (w/v) skim milk and probed with the respective primary (anti-FtsZ, Agrisera. Anti-FtsN [40]. Anti-MinD antisera, gift from Dr. Yu-ling Shih, Academia Sinica, Taiwan). All primary antibodies were used at 1:3333 and secondary antibody (anti-rabbit HRP, 1:10000, Bio-Rad) before being washed and imaged (GE Amersham Imager 600).

## Data availability

All data are presented in the main figures and in the supplementary information.

## Code availability

No new code was generated in this study.

## Acknowledgments

Work in the Söderström lab is funded by the Australian Research Council through an ARC Discovery Project grant (DP220101143). B.S. is supported by an ARC Future Fellowship (FT230100062). A.P. acknowledges the UTS Faculty of Science for a seed funding grant. The authors further wish to acknowledge the use of Nikon N-STORM and Leica Stellaris 8 confocal microscopes within the Microbial Imaging Facility in the Faculty of Science at the University of Technology Sydney.

## Author Contributions

A.P and B.S conceptualized the study. A.P, A.C and B.S. performed the experiments and analysed the data with input from I.D. A.C and M.P contributed reagents. A.P and B.S wrote the manuscript, with editorial input from all authors. All authors approved the final version of the manuscript.

