## Supplementary material for "*E. coli* division machinery drives cocci development inside host cells": Supple

Pokhrel et al.

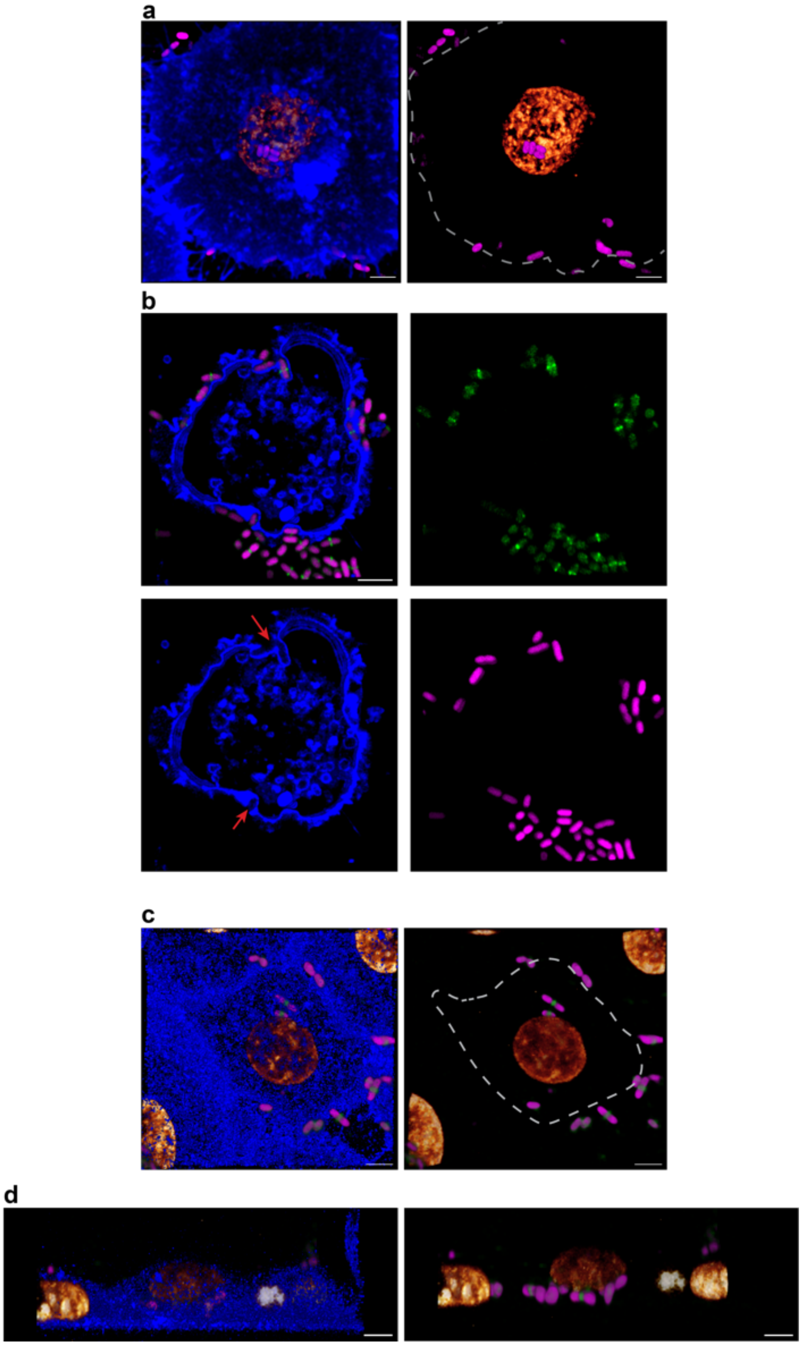


Supplementary figure 1. Infected PD07i bladder cells imaged by confocal microscopy. Nucleus (gold) and membrane (blue). Right side shows images without the membrane channel for better visibility of intracellular bacteria.

**a,** A bladder cell that has taken up UTI89 (Cytoplasmic mCherry: magenta). **b**, A bladder cell in the process of taking up UPEC, red arrows (Cytoplasmic mCherry: magenta, FtsZ-mCitrine: green). **c – d**, A cell where some UTI89 have been taken up (bacterial cells dimmed by the membrane), while others that are external (cells not dimmed by the membrane). Scale bars 4 μm.

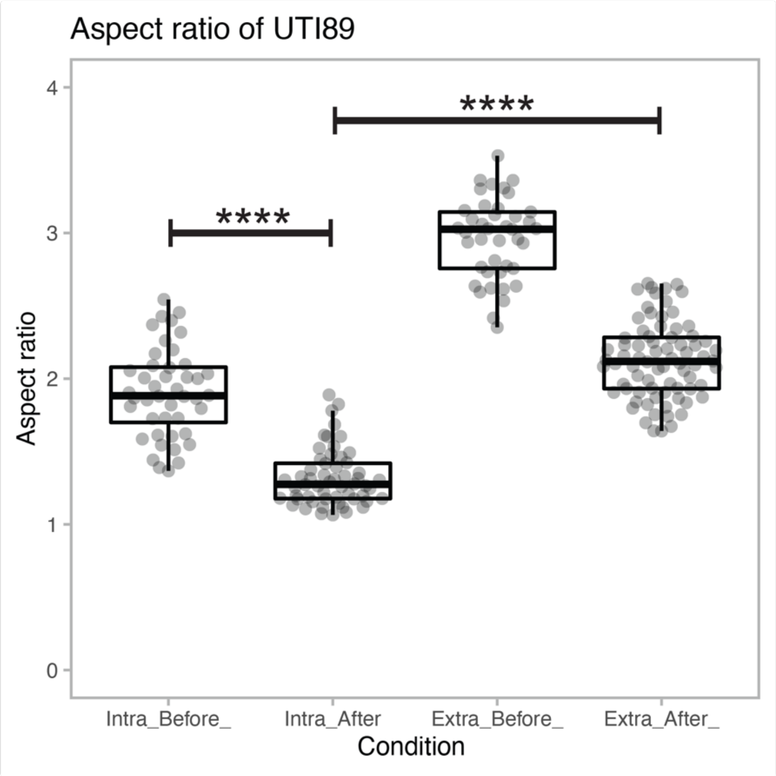


Supplementary figure 2. Aspect ratio (length/width) of intracellular UTI89(mCherry) within PD07i bladder cells, and extracellular UTI89 in EPILIFE growth media, before and after divisions. Four **** (p > 0.0001) indicate extremely statistically significant difference. “Intra” = intracellular bacteria, “Extra” = extracellular bacteria.
Box plots: midline indicates mean, and box represents S.D. Whiskers encompass 1-99% interval of the data.

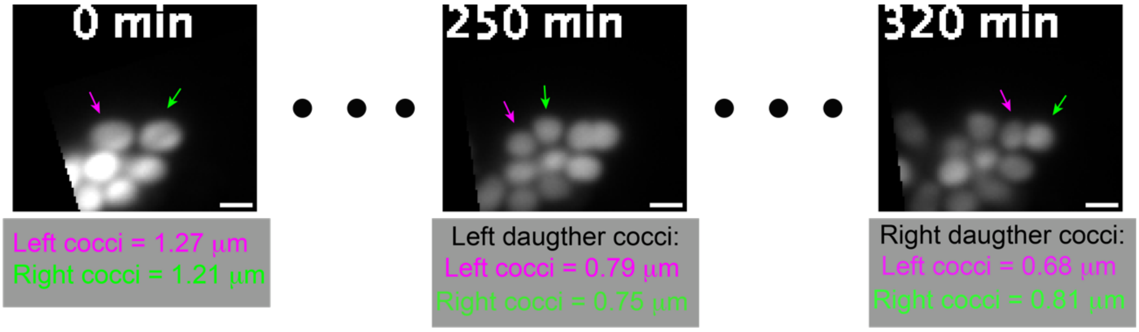


Supplementary figure 3. Still images from a time-lapse image sequence (Supplementary movie SM2) showing UTI89 cocci-to-cocci divisions inside host bladder cells. The UTI89 strain expresses cytoplasmic mCherry (pGI6), pseudocoloured grey. Scale bar 1 μm.

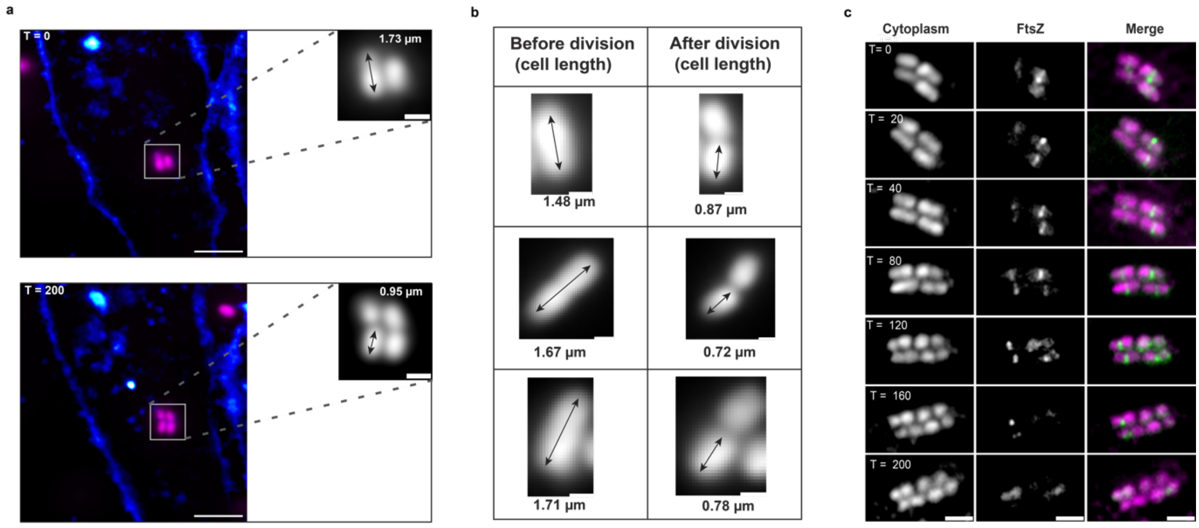


Supplementary figure 4. Clinical UTI isolate MS2027 follows the same rod-to-cocci division patterns inside host cells as UTI89.

**a**, Representative images from a time-lapse sequence of PD07i human epithelial bladder cells (membranes in blue) challenged with MS2027 expressing mCherry (pseudocolored magenta) in the cytoplasm; cells before (T = 0) and after division (T = 200). **b**, Images of intracellular MS2027 cells before and after division across different PD07i human epithelial bladder cells from three separate infections. Example cell lengths of intracellular MS2027 cells before (rods) and after division (cocci) are noted in the table. **c**, Still images from a time-lapse sequence of intracellular MS2027 cells expressing FtsZ-mCitrine, and mCherry in the cytoplasm. Times (T) are shown in minutes. Scale bars: **a** = 5 µm (insets 1µm), **b** = 0.5 µm and **c** = 2 µm.

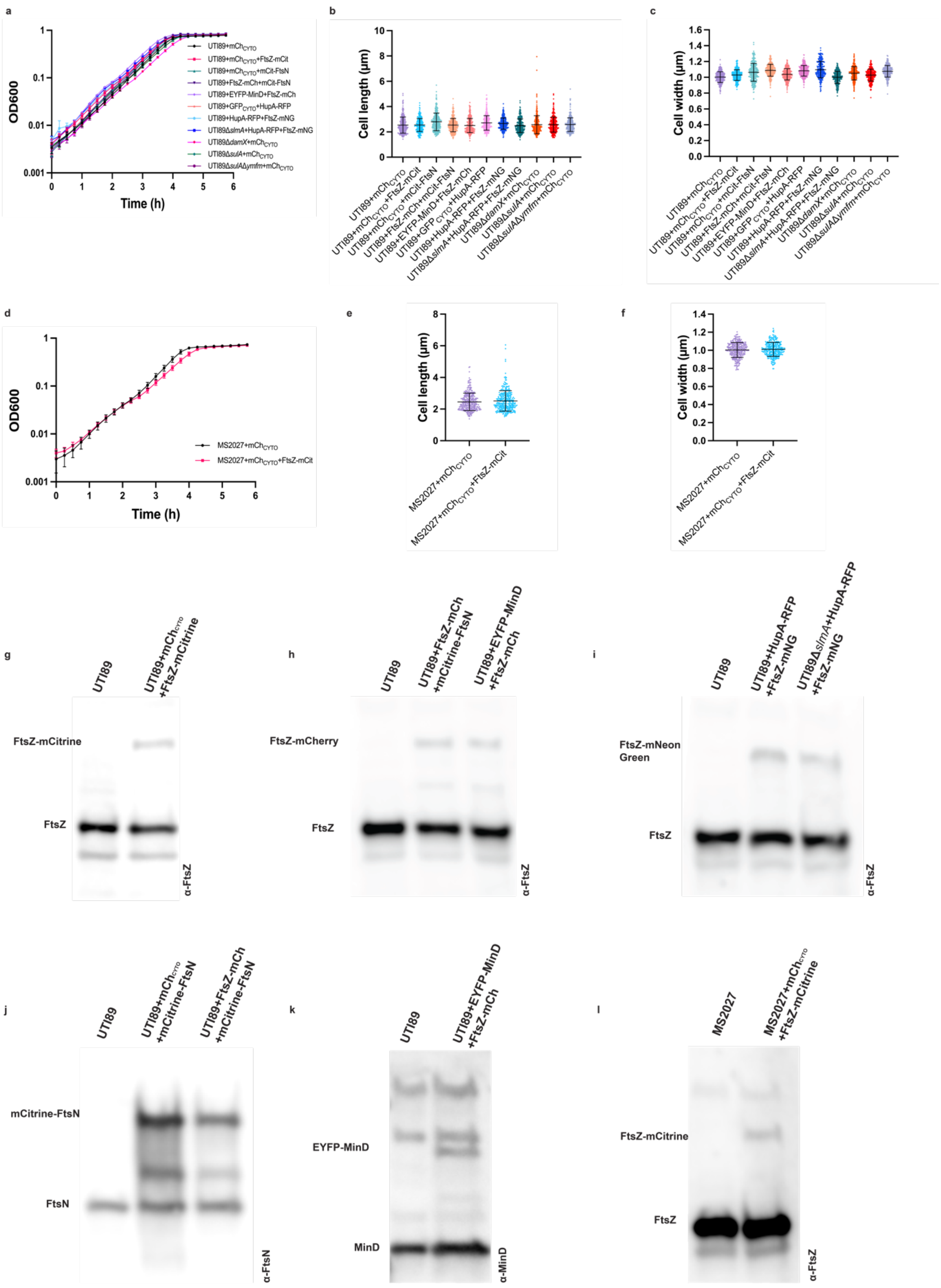


Supplementary figure 5. Bacterial strain viability and quantification of fluorescent protein fusion expression levels.

**a,** Growth curves of UTI89 strains used in the study over a 6h growth period in EPILIFE media. **b - c**, Average cell lengths **(b)** and widths **(c)** of UTI89 strains grown in EPILIFE media in petri-dishes. **d,** Growth curves of MS2027 strains used in the study over a 6h period in EPILIFE media. **e - f**, Average cell length **(e)** and width **(f)** of MS2027 strains grown in EPILIFE media in petri-dishes. All growth curves include data from 3 biological replicates and $n_{cell}$ = 300 for lengths and widths measurements. Midline represents average, and whiskers indicate S.D. **g - l,** Quantitative western blots of FtsZ-mCitrine, FtsZ-mCherry, FtsZ-mNeonGreen, mCitrine-FtsN, and EYFP-MinD production levels in UTI89 and MS2027 cells. FtsZ-mCitrine in **(g)** UTI89, and in **(l)** MS2027 was produced at 13.5(±5.8) % and 7.2(±3.1) % of the total cellular FtsZ, respectively.

**(h)** FtsZ-mCherry in the UTI89+mCitrine-FtsN and in UTI89+EYFP-MinD strains was produced at 16.8(±8.3) % and 18.5(±7.9) % of the total cellular FtsZ respectively. **(i)** FtsZ-mNeonGreen in the UTI89+HupA-RFP and in UTI89Δ*slmA* +HupA-RFP strains was produced to 18(±2.6) % and 17.4(±3.1) % of the total FtsZ.

**(j)** mCitrine-FtsN in UTI89+$mCh_{CYTO}$ and UTI89+FtsZ-mCherry strains was produced to approximately 142(±14.5) % and 118.7(±1.5) % of the native FtsN production, respectively. Overall, these values for FP-FtsN and FtsZ-FP have previously been shown not interfere with growth and division during vegetative growth in rich media [1-3]. **(k)** EYFP-MinD production in the UTI89+FtsZ-mCherry strain showed that EYFP-MinD was produced at approximately 28(±6.4) % of the total cellular MinD. Note some unspecific binding of the anti-MinD antisera. Collectively, these results show that the fluorescent protein fusions were not heavily overexpressed in any of the strains used in this study. All percentages (± S.d.) are based on three independent experiments. mCh = mCherry, mCit = mCitrine, mNG = mNeonGreen, $GFP_{CYTO}$ = Free GFP in the cytoplasm, $mCh_{CYTO}$ = Free mCherry in the cytoplasm.

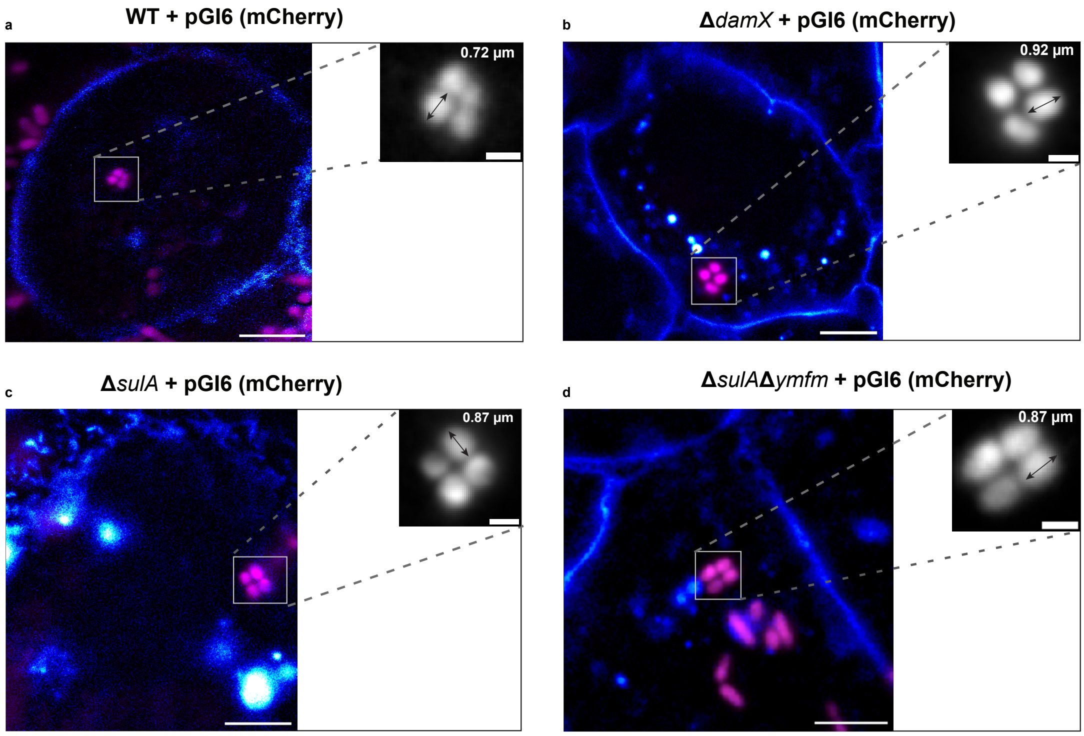


Supplementary figure 6: Deletion of genes known to regulate UPEC cell morphology and division during late stages of infection does not affect rod-to-cocci divisions. Representative images of PD07i human epithelial bladder cells (membrane in blue) challenged with **a,** WT UTI89, **b,** UTI89Δ*damX,* **c,** UTI89Δ*sulA* and **d**, UTI89Δ*sulA*Δ*ymfM.* All strains expressed mCherry from pGI6 in the cytoplasm as a volume marker (pseudo-coloured magenta). Example cocci lengths of parental UTI89 and mutant strains after division are noted in the insets. Scale bars **a - d** = 5 μm (insets 1μm).

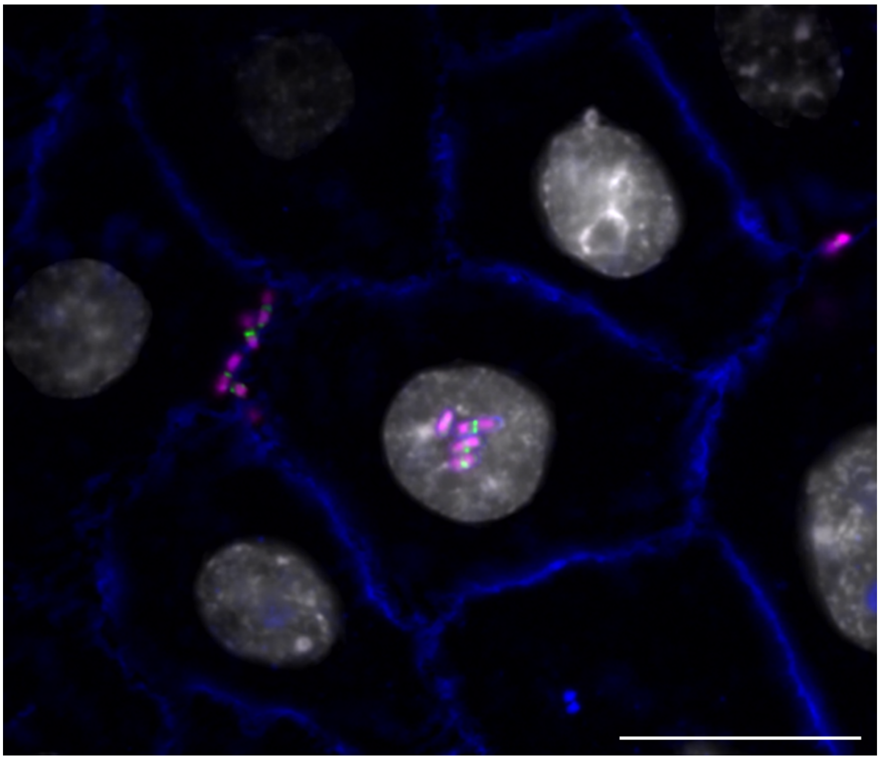

Supplementary figure 7: Representative HILO microscopy image of PD07i bladder cells challenged with UTI89/HupA-RFP/FtsZ-mNeonGreen. PD07i membrane in blue, nucleus in grey, HupA in magenta and FtsZ in green. Scale bar 20 μm.

*Table 1: Bacterial strains and plasmids used in the study.*

| Strain | Plasmid(s) | Fluorescent protein(s) | Inducer | Source | Reference to Figure |
|---|---|---|---|---|---|
| UTI89 | pGI6 | $mCherry^{CYTO}$ | - | [4] | Fig.1, Supp. Fig 1, Supp. Fig 3, Supp. Fig 6 |
| UTI89 | pGI6 / pHC054 | $mCherry^{CYTO}$/ FtsZ-mCitrine | IPTG | [4,5] | Fig., Supp. Fig 1 |
| UTI89 | pGI6 / pHC004 | $mCherry^{CYTO}$/ mCitrine-FtsN | Rhamnose | [2,4] | Fig.1 |
| UTI89 | pMP11 / pHC004 | FtsZ-mCherry/ mCitrine-FtsN | Rhamnose | This Study/[2] | Fig.1 |
| UTI89 | pSR-4 | EYFP-MinD | IPTG | [6] | Fig.2 |
| UTI89 | pMP11 / pSR-4 | FtsZ-mCherry/ EYFP-MinD | IPTG | This Study /[6] | Fig.2 |
| UTI89 | pGI5 / pSTC011 | $msfGFP^{CYTO}$/ HupA-RFP | IPTG | [7,8] | Fig.2 |
| UTI89 | pMP7 / pSTC011 | FtsZ-mNeonGreen/ HupA-RFP | IPTG | This Study/[8] | Fig.2, Supp. Fig 7 |
| UTI89Δ*slmA* | pMP7 / pSTC011 | FtsZ-mNeonGreen/ HupA-RFP | IPTG | This Study/[8] | Fig.2 |
| UTI89Δ*sulA* | pGI6 | $mCherry^{CYTO}$ | - | [4,9] | Supp. Fig 6 |
| UTI89Δ*sulA* Δ*ymfm* | pGI6 | $mCherry^{CYTO}$ | - | [4,10] | Supp. Fig 6 |
| UTI89Δ*damx* | pGI6 | $mCherry^{CYTO}$ | - | [4,11] | Supp. Fig 6 |
| MS2027 | pGI6 | $mCherry^{CYTO}$ | - | [4,12] | Supp. Fig 4 |
| MS2027 | pGI6 / pHC054 | $mCherry^{CYTO}$/ FtsZ-mCitrine | - | [4,5,12] | Supp. Fig 4 |

Plasmid engineering

To engineer pMP7, the *ftsZ-mNG* sandwich fusion, originally sourced from BW27783/FtsZ-55mNeonGreen56 [13] (gift from Harold Erickson, Duke University), was PCR amplified from plasmid pDD3 (gift from Daniel Daley, Stockholm University) using primers 1 and 2. The vector backbone, pGI3 [7], was digested with *NcoI* and *BamHI*. Both DNA fragments were then purified and ligated by in vivo DNA synthesis. To make pMP11, *mCherry* was PCR amplified from pDSW931[14] (Gift from David Weiss, University of Iowa) using primers 3 and 4, and the vector, pMP7, was amplified using primers 5 and 6. Both DNA fragments were then purified and ligated by in vivo DNA synthesis. The sequences were verified by Sanger sequencing (AGRF, Australia).

Primers:

1. GATAGCGCCCGGTCTAGAGGAGGTACTACCATGTTTGAACCAATGGAACTTACC
2. GGCTGCAGGTCGACGGATCTTTAGGATCCTTAATCAGCTTGCTTACGCAGG
3. GGTTGGAGGATCCACCCTCGAGATGGTGAGCAAGGGCGAGGAG
4. GAATCGTCTGGGTGGATCCCTCGAGCTTGTACAGCTCGTCCATGCCG
5. CTCGAGGGATCCACCCAGACGATTCAAATC
6. CATCTCGAGGGTGGATCCTCCAACCG

Supplementary Movie SM1
Time-lapse of a bladder cell (membrane in grey) invaded by UTI89 (mCherry pseudo-coloured magenta). Gentamicin protection assay, f.c. 50 μg $ml^{-1}$. Note that only intracellular bacteria are viable. Imaged using HILO illumination. Scale bar 2 μm.

Supplementary Movie SM2
Time-lapse of UTI89 cocci to cocci divisions (mCherry pseudo-coloured grey). Imaged using HILO illumination. Scale bar 1 μm.

Supplementary Movie SM3
Time-lapse of intracellular UTI89 (mCherry pseudo-coloured magenta) expressing FtsZ-mCitrine (green). Note at T= 90 min that FtsZ-mCitrine rings have disassembled. Imaged using HILO illumination. Scale bar 1 μm.

Supplementary Movie SM4
Time-lapse of intracellular UTI89 (mCherry pseudo-coloured magenta) expressing mCitrine-FtsN (green). Imaged using HILO illumination. Scale bar 1 μm.

Supplementary Movie SM5
Time-lapse of intracellular UTI89 expressing FtsZ-mCherry (left) and mCitrine-FtsN (mid). Merged channels (right) FtsZ-mCherry (magenta) and mCitrine-FtsN (green). Imaged using HILO illumination. Scale bar 1 μm.

Supplementary Movie SM6
Time-lapse of intracellular UTI89 expressing EYFP-MinD. Imaged using HILO illumination. Scale bar 1 μm.

Supplementary Movie SM7
Time-lapse of extracellular UTI89 expressing HupA-RFP (left) and FtsZ-mNeonGreen (mid) and merged (HupA-RFP in magenta, mNeonGreen in green). Prior to addition of Gentamicin. Imaged using HILO illumination. Scale bar 4 μm.

Supplementary Movie SM8
Time-lapse of extracellular UTI89 expressing HupA-RFP (magenta) and FtsZ-mNeonGreen (green) with 50 μg $ml^{-1}$ Gentamicin added to the EpiLife growth media. The chromosomes are condensed to midcell and FtsZ rings are disassembled. Time between Gentamicin addition to media in the dish and imaging < 10 minutes. Imaged using HILO illumination. Scale bar 4 μm.
